# Structural Basis of *Pseudomonas* Biofilm-Forming Functional Amyloid FapC Formation

**DOI:** 10.1101/2025.03.15.642095

**Authors:** Kasper Holst Hansen, Mert Golcuk, Chang Hyeock Byeon, Abdulkadir Tunc, Emilie Buhl Plechinger, James F. Conway, Maria Andreasen, Mert Gur, Ümit Akbey

## Abstract

Biofilm-protected *Pseudomonas aeruginosa* causes chronic infections that are difficult to treat. FapC, the major biofilm forming functional-amyloid in *Pseudomonas*, is essential for biofilm integrity, yet its structural details remain unresolved. Using an integrative structural biology approach, we combine solution NMR-based structural ensemble of unfolded monomeric FapC, a ∼3.3 Å resolution CryoEM density map of FapC fibril, and all-atom MD simulations to capture transition from unfolded to folded monomer to fibrillar fold, providing a complete structural view of FapC biogenesis. CryoEM reveals a unique triple-layer β-solenoid cross-β fibril composed of a single protofilament. MD simulations initiated from monomeric and fibrillar FapC mapped structural transitions, offering mechanistic insights into amyloid assembly and disassembly. Understanding FapC reveals how *Pseudomonas* exploits functional amyloids for biofilm formation and establishes a structural and mechanistic foundation for developing therapeutics targeting biofilm-related infection and antimicrobial resistance.

## 1. Introduction

Amyloids are found across diverse organisms, from bacteria to humans, and are commonly associated with protein misfolding and aggregation into fibrils linked to neurodegenerative diseases.^1,2^ A subgroup of amyloids, functional bacterial amyloids (FuBAs), in contrast plays essential biological roles, particularly in biofilm formation.^3^ Biofilm-forming functional amyloids (FAs) serve as key structural components within microbial communities, embedded in extracellular matrix polymers composed of polysaccharides, amyloid proteins, and extracellular DNA (eDNA).^4^ Biofilm-producing bacteria are a significant health concern since 80% of all chronic human infections are either related to or caused by both biofilm formation and biofilm-mediated resistance to antimicrobial and antibiotic therapies.^5^ Fighting biofilm protected bacteria requires detailed structural information of biofilm components for a directed targeting by pharmaceutics. Yet such structural information remains scarce, creating a critical gap that hinders the development of targeted therapeutics in chronic biofilm-associated infection treatments. High-resolution structural determination of functional amyloid fibrils in bacterial biofilms represents a promising approach to design anti-biofilm agents, disrupt biofilms, and improve treatments for chronic infections.

The biofilm forming Gram-negative *Pseudomonas aeruginosa* is recognized by the World Health Organization (WHO) as one of the 15 most critical human pathogens, responsible for chronic infections such as those seen in cystic fibrosis.^6,7^ In *Pseudomonas*, the Fap (Functional Amyloid in Pseudomonas) system plays a central role in biofilm formation and is tightly regulated. This system is encoded by the Fap operon, which consists of six proteins: FapA– F.^8,9^ FapA functions as a translational regulator of FapAB, and previous studies—including our own—suggest that FapA may also act as a chaperone or chaperone-like protein that influences amyloid formation.^9-12^ The main extracellular FA secreted in the *Pseudomonas* biofilm are FapB and FapC, with FapC being the predominant component. Both proteins are intrinsically disordered as monomers, and FapB serves as a nucleator for FapC polymerization.^8,9^ Moreover, FapE has been detected in native Fap fibrils.^9^ Upon secretion into the biofilm environment, these proteins assemble into highly ordered cross-β amyloid filaments, forming the structural scaffold of biofilms. Other Fap system components include FapD, a protease that modifies Fap proteins, and FapF, a trimeric membrane transporter responsible for secreting Fap proteins into the biofilm matrix.

In recent years, we and others have characterized the biophysical and kinetic properties of FapC.^8,9,13-19^ However, structural studies have been limited in resolution, leaving detailed structures of FapC monomers and fibrils unknown. Using solution Nuclear Magnetic Resonance (NMR), we previously provided complete resonance assignment of FapC and FapA from *P. aeruginosa*, identifying secondary structure propensities (SSPs) for these intrinsically disordered proteins (IDPs) and characterizing their interactions.^3,12,19,20^ However, no high-resolution structure of FapC has been determined to date, either in its monomeric or fibrillar state. The only available structural models include one derived from sequence covariance data^21^ and an AlphaFold2 (AF) AI/ML-generated monomeric model (AF-C4IN70). However, nearly half of the residues in the AF model are predicted with low or very low confidence, as indicated by the predicted Local Distance Difference Test (pLDDT) scores: 6.8% Very High Confidence (>90), 55.2% High Confidence (70–90), 20% Low Confidence (50–70), and 18% Very Low Confidence (≤50). Furthermore, because the AF model represents a single FapC monomer, it does not account for inter-subunit interactions essential for fibril formation and stability.

Here, we present the first high-resolution structural description of both the monomeric and fibrillar form of FapC from *Pseudomonas* derived from *in vitro* samples. Using solution NMR, we characterize the ensemble conformations of FapC in its unstructured monomeric state. To resolve the fibrillar structure, we employed cryogenic electron microscopy (cryo-EM) to generate a high-resolution density map and used the AF model as a starting reference for developing a refined fibrillar FapC model. This cryo-EM-based structure was further validated and refined using all-atom molecular dynamics (MD) simulations, which assess structural stability and investigate the role of protein-protein interactions within the fibril, particularly between neighboring FapC subunits. Our MD simulations reveal that the AF model aligns more closely with our MD-optimized monomeric cryo-EM fibril model, whereas in the fibrillar state (simulated as a trimer), the MD simulation stabilizes at around our cryo-EM-derived fibril structure. By combining conventional and steered MD (SMD) simulations, we characterized intermediate conformational states and transitions, bridging the gap between the experimentally determined unfolded monomer and the fibrillar structure of FapC. Altogether, our findings provide a comprehensive model for the full biogenesis of FapC, covering its unstructured monomer, structured monomer, fibrillar state, and intermediate conformations. Understanding the structural basis of FapC functional amyloid formation is a crucial step toward deciphering *Pseudomonas* biofilm biogenesis and will ultimately facilitate the rational design of amyloid-modifying agents to combat persistent *Pseudomonas* infections.

## 2. Methods

### FapC preparation

Bacterial plasmids were transformed into the BL21(DE3) E. coli bacteria containing the recombinant FapC residues between 25-250. The first 24 aminoacids forms the signal sequence that is cleaved upon secretion and not part of the functional fibril in vivo.^21^ All unlabeled samples were prepared in LB media. Isotope labeling for NMR studies was carried out by growing in minimal media ^15^N-ammonium chloride (1 g/L) as the nitrogen source and ^13^C-D-glucose (2 g/L). The glycerol stocks were grown overnight in LB media. The overnight culture cells were harvested and resuspended in the large growth media with appropriate antibiotic. These cultures were grown in shaker incubator at 37 °C until the OD600 reached a value between 0.8-1.0 OD. The cultures were induced with 1 mM IPTG final concentration and grown for another 4 h at 37 °C. Cells were harvested by centrifugation (6000 RCF for 20 min at 4 °C). The cells were then lysed using a sonicator in Lysis buffer (50 mM Tris, pH 8 and 8 M guanidinium chloride (GuHCl), then the soluble lysate was separated from the insoluble cell debris using centrifugation (6000 RCF for 20min at 4 °C). The purification was done by using HisPur Ni-NTA resin (Thermo Scientific) by step-eluting the N-terminally His-tagged FapC in imidazole containing Lysis buffer. The MW of FapC is 23.755 kDa. The soluble lysate solution was incubated overnight at 4 °C with Ni-NTA resin. The resin was separated by centrifugation (6000 RCF for 20 min at 20 °C) and the supernatant was decanted off. The resin was washed with 35 mL of Lysis buffer and centrifuged to separate the resin. Additional washes and step elutions were done in a similar manner following this order: 35 mL wash with Lysis buffer, 25 mL elutions of 30, 60, 120, 300 and 500 mM imidazole buffers. Elutions were pooled and concentrated using a spin filter concentrator with 3000 MW-cutoff.

### Expression and purification of native *Pseudomonas* sp. UK4 Fap fibrils

Electrocompetent UK4 Δ*fap* cells were prepared from 6 mL of overnight culture grown in LB medium at 30°C.^22^ Briefly, cells were harvested by centrifugation (7500g, 10 min), washed twice in 4 mL of 300 mM room temperature sucrose, and finally suspended in 100 μL of 300 mM sucrose. 5 μL of pMMB190Tc-UK4fap plasmid (∼400 ng/μL) added to the electrocompetent cells and 40 μL of the suspension was transferred to a 1-mm gap electroporation cuvette.^10^ Electroporation was performed with a Micro Pulser electroporator (Biorad, Hercules, CA) using a 1.25 kV pulses. 1 mL LB medium was applied directly after electroporation and the sample was transferred to a 15-mL tube and incubated (28°C, 200 rpm, 2 h). 100 μL of the transformations were plated on LB agar plates containing 50 μg/mL tetracycline and the plates were incubated at 30°C until visible colonies appeared (1–3 days). A colony of the transformed cells was grown over night in 10 mL LB medium containing 50 μg/mL tetracycline, and 100 μL of the overnight culture was spread on a nematode growth medium (NGM) agar plate (2.5 g/L peptone, 3.0 g/L NaCl, 5 mg/L cholesterol, 17.5 g/L agar) with 50 μg/mL tetracycline. NGM medium was used as we have found it reduces production of contaminating exopolysaccharides compared to LB and CFA medium (unpublished results). The plate was incubated (28°C, 3 days), after which biomass was harvested by scraping the plate using a cell scraper and transferred to a 2 mL microcentrifuge tube with screw cap for amyloid purification. The harvested biomass was suspended in 900 μL of buffer (10 mM Tris-HCl, pH 8.0) and homogenized using a pipette. 300 μL of enzyme mix (0.4mg/mL RNase A, 0.4mg/mL DNase I, 4mg/mL lysozyme, 4mM MgCl_2_ and 0.4% Triton X-100) was added to the suspended cells, and the cells were lysed by three times freeze thawing using a -80°C freezer and a water bath at 37°C. The suspension was then incubated (37°C, 2hr, without shaking) to allow the enzymes to digest the peptidoglycan and nucleic acids. After the incubation,133 μL of 20% SDS was added and the sample boiled for 5 min in a water bath to unfold and solubilize non-amyloid proteins. The sample was centrifuged (16,000x g, 10min) and the supernatant discarded. The pellet was resuspended in 1200 μL buffer, after which 133 μL of 20% SDS was added and the heating a centrifugation repeated. The sample was hereafter washed twice with 1.2 mL buffer by centrifugation and resuspension and finally resuspended in 250 μL buffer.

### Aggregation kinetics

Samples in 8 M GuHCl were desalted using PD Minitrap G-10 columns (Cytiva) into 20 mM Sodium Phosphate pH 7.8, 0.02% w/v Sodium Azide. Sample concentrations were measured using UV280. FapC concentration was adjusted to 50 µM with 20 mM Sodium Phosphate pH=7.8, 0.02% w/v/ Sodium Azide and Thioflavin T (ThT) was added to a final concentration of 200 mM. 50 µL volumes of samples were loaded into Corning 3881 (Sigma Aldrich) plates. The ThT fluorescence response was measured at a Tecan Spark at 442/482 nm excitation/emission every 10 min for 48 hours at 37 °C. All measurements were done as triplicates.

### Synchrotron Radiation Circular Dichroism (SR-CD)

The SR-CD spectra of the FapC aggregates and monomers were collected at the AU-CD beamline of the ASTRID2 synchrotron, Aarhus University, Denmark. Aggregated FapC was centrifuged (13,000 rpm for 60 min), supernatants discarded, and the pellet resuspended in the same volume of 20 mM phosphate buffer, pH 7 and sonicated for 2 s using a probe sonicator. Following purification and desalting the monomer sample was desalted a second time using a PD-10 column to remove additional guanidinium chloride. Three to five successive scans over the wavelength range from 180 to 280 nm were recorded at 25°C, using a 0.2 mm path length cuvette, at 1 nm intervals with a dwell time of 2 s. All SR-CD spectra were processed and subtracted from their respective averaged baseline (solution containing all components of the sample, except the protein), smoothing with a seven pt Savitzky-Golay filter, and expressing the final SRCD spectra in mean residual ellipticity. The SR-CD spectra were deconvoluted using DichroWeb,^23,24^ to obtain the contribution from individual structural components. Each spectrum was fitted using Selecon3, Contin, and CDSSTR programs with the SP180 reference data set,^25^ and an average of the structural component contributions from the three analysis programs were used.

### Nuclear Magnetic Resonance (NMR)

NMR spectra of uniformly ^13^C,^15^N isotope labeled FapC FL (FapC_25-250_: 226 aa, without the signal peptide) were recorded at Bruker Avance III 600, Avance 700 and Avance II 900 MHz spectrometers equipped with 5 mm triple resonance TCI cryoprobes. These experiments were used to assign FapC as previously published.^19^ Samples of protein concentration ∼0.3mM in 50 mM sodium phosphate, 30 mM DTT, 1 mM D_6_-DSS, 10% D_2_O and pH 7.4 buffer were placed in 5 mm NMR sample tubes with a total volume of 500 μl. Samples were prepared freshly prior to performing the NMR experiment. The following experiments were used in accomplishing the full assignment: 2D HN HSQC and 3D HNCACB, HN(CO)CACB, CC(CO)NH TOCSY, H(CCCO)NH TOCSY and HBHA(CO)NH. The 3D 13C-edited/15N-edited NOESY HSQC experiment was used to further confirm the assignment. In this paper, further analysis of the NOSEY experiment was done to generate NOE distance restraints for structure calculations. All spectra except 3D NOESEY were processed using Topspin 3 (Bruker Biospin). NMRPipe was used to process the time-shared 3D NOESY experiment NMRPipe.^26^ The NOE restraints were manually generating with the use of CCPNmr Analysis 3.^27^ The ^1^H chemical shifts were referenced to 0 ppm by using DSS as an internal standard added to the NMR samples. The ^13^C and ^15^N chemical shifts were indirectly referenced by using the ^1^H frequency as explained previously.^28^

Solid-state NMR (ssNMR) experiments were performed on a Bruker Avance III 750 spectrometer equipped with a 3.2 mm triple resonance HCN LT MAS probe. The homonuclear 2D ^13^C-^13^C correlation spectra were recorded with 20 ms DARR or 10 ms TOBSY polarization mixing schemes.^29,30^ 32 scans were recorded for 512 increments with 1-2 second recycle delay. SPINAL64 heteronuclear decoupling was applied using ∼90 kHz RF strengths on protons.^31^ 2 ms contact time was used for the CP spectrum. All spectra were processed using Topspin 3.6 (Bruker Biospin). **Monomeric structural ensemble calculation by solution NMR**. From the NOESY experiments, 706 total NOE restrains were manually generated. With the chemical shifts, TALOS was used to generate 352 dihedral angles.^32^ The anneal.py protocol in XPLOR-NIH was used to perform the structure calculations with the NOE and dihedral angle restrains.^33^ ∼10k structures were generated from this calculation with low violations. Chemical shift assignments can be found in the Biological Magnetic Resonance Data Bank for FapC FL (#51793).

### Negative-stain Electron-Microscopy sample preparation and data collection

400 mesh copper grids with a continuous carbon film were glow discharged for 90 seconds. ∼3 µL of FapC was applied to grids and left for 10 seconds before side blotting on filter paper. Grids were stained with 2% (weight/volume) uranyl acetate for 10 seconds before side blotting on filter paper. Micrographs were recorded on a Tecnai TF20 microscope (Thermo Fisher Scientific - TFS, MA, USA) with a field emission gun operating at 200 kV equipped with a TVIPS XF416 CMOS camera (TVIPS GmbH, Gilching, Germany). No further processing was done to the micrographs shown.

### CryoEM sample preparation and data collection

Quantifoil R2/1 Cu 300 mesh holey carbon grids (Protochips, Morrisville, NC, USA) were glow discharged for 45 s at 25 mA using an EmiTech K100X (Quorom Technologies Ltd, Laughton, ES, UK). 3 μL of FapC (0.5 mg/mL) was applied to grids, blotted for 6 s, and plunge-frozen in liquid ethane using a Vitrobot Mark IV plunge freezer with a blot force of zero (TFS) maintaining 4°C and 99% humidity. Vitrified grids were stored in liquid nitrogen until imaging. Data collection was performed at the University of Pittsburgh Cryo-EM Center. A Titan Krios G3i was utilized operating at 300 kV with a Selectris energy filter and Falcon 4i camera under control of the EPU v3.2 software (TFS). The nominal magnification was 165,000x (pixel size of 0.72 Å) and images were collected with the energy filter slit width set to 10 eV and under-focus ranging between -0.5 and -2.0 μm. Exposures were 4.8 seconds for a total dose of ∼45 electrons/Å^2^. A total of 8870 movies were collected as movies in EER format.

### Helical reconstruction

All data processing was performed in CryoSPARC version 4.5.^34^ Movies were drift corrected using Patch Motion Correction. Patch CTF was used to estimate Contrast Transfer Function (CTF) parameters. Particle picking was performed with Filament Tracer using template free mode with estimates of fibril dimensions from AF and prior refinement work on FapC. ∼762k particles were extracted with a box size of 300 pixels and then manually curated to a resolution estimate of 6 Å or better for further analysis. At this point, only one type of 2D Class was observed, indicating the presence of a single type of FapC filament. A single round of reference-free two-dimensional (2D) classification was used to remove the noisiest particles. To remove further noise from the particle set, two rounds of 2D classification and 6 rounds of 3D classification were performed. 3D classification was run in simple initialization mode using a solvent mask generated from an initial helical reconstruction. Particles were then reconstructed using helical refinement with symmetry estimates of -2.5° twist, 14.7 Å rise, and C1 symmetry. Afterwards, particles were CTF refined using global- and local CTF refine. Particles were reconstructed once more using the same symmetry parameters. Finally, a single round of 2D classification was used to remove the final bit of noise from the particle set. The now ∼72k particles were reconstructed into the final density map using helical refinement. The final map had a Rise of 14.562 Å and a twist of -2.266°. Overall resolution estimated by CryoSPARC was ∼3.3 Å according to the gold standard Fourier shell correlation (GSFSC) cutoff at 0.143. The final map was sharpened by using DeepEMhancer.^35^

### Model building and validation

Model building was performed with UCSF ChimeraX 1.8^36^ using the real-time MD software package ISOLDE.^37^ The AF FapC model was utilized as a starting point due to the overlap of the predicted model to the Cryo-EM density. To build the FapC structural model we utilized both CryoSPARC density map and DeepEM tight target map. The later represents much better resolution particularly with regards to sidechain densities, however, has weaker densities at some of the b-strand regions. The model was improved by iterative fitting in ISOLDE to fix clashes and unfavored conformations. Finally, segments of β-strands favoring sequences were identified and the direction of backbone was manually adjusted to facilitate β-sheet hydrogen bonding. The final model was validated by PHENIX.^37,38^ The final FapC fibril model has excellent Ramachandran scores with 97% favored and 3% allowed orientations were obtained with low clash/MolProbity scores with a high density to model overlap. See **Table SI1** for details. CryoEM density was deposited to the EMDB with accession code (**EMD-49649**). The final FapC model was deposited to the PDB with accession code (**9NQD**).

### Molecular Dynamics simulations

The atomic models of FapC, derived from both NMR data and AF predictions, were placed in a cubic box of TIP3P water with dimensions of 160 Å along each Cartesian axis. The systems were neutralized and brought to a physiological ionic concentration of 150 mM KCl, resulting in a total of approximately 400,000 atoms per system. All preparatory steps, including solvation and ionization, were performed using VMD. MD simulations were performed in NAMD 3,^39^ with the CHARMM36 all-atom force field,^40^ using a 2 fs time step. The temperature was maintained at 310 K by Langevin dynamics with a damping coefficient of 1 ps^−1^, and the pressure was kept at 1 atm using the Langevin Nosé–Hoover method (oscillation period of 100 fs, damping timescale of 50 fs). A 12 Å cut-off was employed for van der Waals interactions, and long-range electrostatic interactions were computed by the particle-mesh Ewald (PME) method. The systems underwent two rounds of energy minimization and equilibration. In the first round, the protein coordinates were fixed while the system was minimized for 10,000 steps, followed by a 2 ns equilibration to allow the solvent to relax around the protein. In the second round, the protein was released, and another 10,000 steps of minimization were performed, followed by a 4 ns stabilization phase under harmonic restraints of 1 kcal·mol^−1^·Å^−2^ on the C_α_ atoms. Afterward, all restraints were removed, and the system was further equilibrated for 4 ns before the production phase. For each atomic model of FapC, three independent MD simulations were performed. During the production phase, coordinates were saved every 0.1 ns and used for all subsequent analyses.

### Steered Molecular Dynamics simulations

SMD is a computational technique in which external force is applied to specific atom(s) to guide conformational changes along a predefined pathway.^41^ FapC conformations were aligned along the x-axis by designating the C_α_ atom of the N-terminal residue (G25) as the fixed point and the C_α_ atom of the C-terminal residue (F250) as the pulling point, thereby defining the vector along which the external force in SMD simulations was applied. To accommodate the extension in the pulling direction, the systems were resolvated in a water box providing 130 Å of padding along the pulling axis and 25 Å of padding in the perpendicular directions. Constant-velocity SMD simulations were performed with a pulling velocity of 0.1 Å/ns and a spring constant of 100 kcal·mol^−1^·Å^−2^, where a dummy atom connected by a spring to the steered atoms was pulled at the specified velocity. After 100 Å of pulling, the fixed and pulling atoms were reassigned, because the original N- and C-termini became unstructured. The new fixed and pulling atoms were set to the N- and C-termini of the structured fold (**Table SI2**), enabling continued simulation of the mechanical unfolding behavior for the remaining structured regions in a controlled manner.

### Principal Component Analysis

Principal Component Analysis (PCA) is a statistical method that identifies dominant patterns of structural fluctuations by reducing the dimensionality of protein conformational ensembles. The FapC structures obtained from the NMR were aligned using the C_α_ atoms. The FapC structures obtained from the NMR ensemble were first superimposed on their C_α_ atoms to minimize the root-mean-square deviation (RMSD), which measures the average atomic displacement between protein structures. Using these aligned coordinates, the covariance matrix was constructed as: **C= ⟨(R − ⟨R⟩)(R − ⟨R⟩)**^**T**^**⟩**.^42^ Here **R** is the 678-dimensional configurational vector, composed of the instantaneous coordinates of the 226 C_α_ atoms of FapC, and **⟨R⟩** is the ensemble average of **R**. The diagonal elements of the covariance matrix **C** (where *i* **=** *j*) represent the atomic mean square fluctuation (MSF), expressed as **⟨(ΔR**_**i**_)^**2**^**⟩**, whose square root corresponds to the root-mean-square fluctuation (RMSF), and both measures indicate the degree of flexibility of individual atoms in the structural (NMR) and conformational (MD) ensembles. **C** was then diagonalized via eigenvalue decomposition to extract the principal components (PCs): 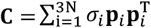. Here, **p**_***i***_ denotes the *i*^th^ PC, and *σ*_*i*_ is the corresponding eigenvalue (variance), which quantifies the magnitude of motion along that PC. The PCs are ranked in descending order based on their variances *σ*_*i*_. Because PC1 and PC2 have the largest variances, they capture the dominant motions in the structural (NMR) and conformational (MD) ensembles. To project each conformation onto the i^th^ PC, the following operation was performed: 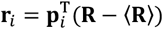. These projections (**r**_*i*_) provide insights into the conformational dynamics of FapC along the *i*^th^ PC.

## 3. Results

### Ensemble Description of Monomeric FapC

The historical domain organization consensus of FapC in UK4 *P. aeruginosa* (NR1L1R2L2R3C, residues 25-250) is based on amino acid composition and repeat occurrence. FapC contains three imperfect repeats (R1-R3, ∼30 residues each), two loops (L1, L2, ∼35/43 residues), and N-/C-terminal regions (∼37/21 residues) (**Fig. 1A, SI1**). The secondary structure elements of the unfolded FapC monomer, derived from our complete resonance assignment, closely match the AF-predicted monomer model and correspond to areas with a high level of confidence in the AF prediction.^19^ However, they deviate significantly from the traditional domain consensus, **Fig. SI1**. To characterize the unfolded monomer, we used high-resolution 3D solution NMR, confirming that FapC behaves as an intrinsically disordered protein (IDP). The well-resolved cross-peaks in the 2D ^1^H-^15^N HSQC spectrum (**Fig. 1B**) are concentrated within a narrow proton chemical shift range, indicative of its IDP nature, shown with selected assignments.^19^

**Figure 1:**
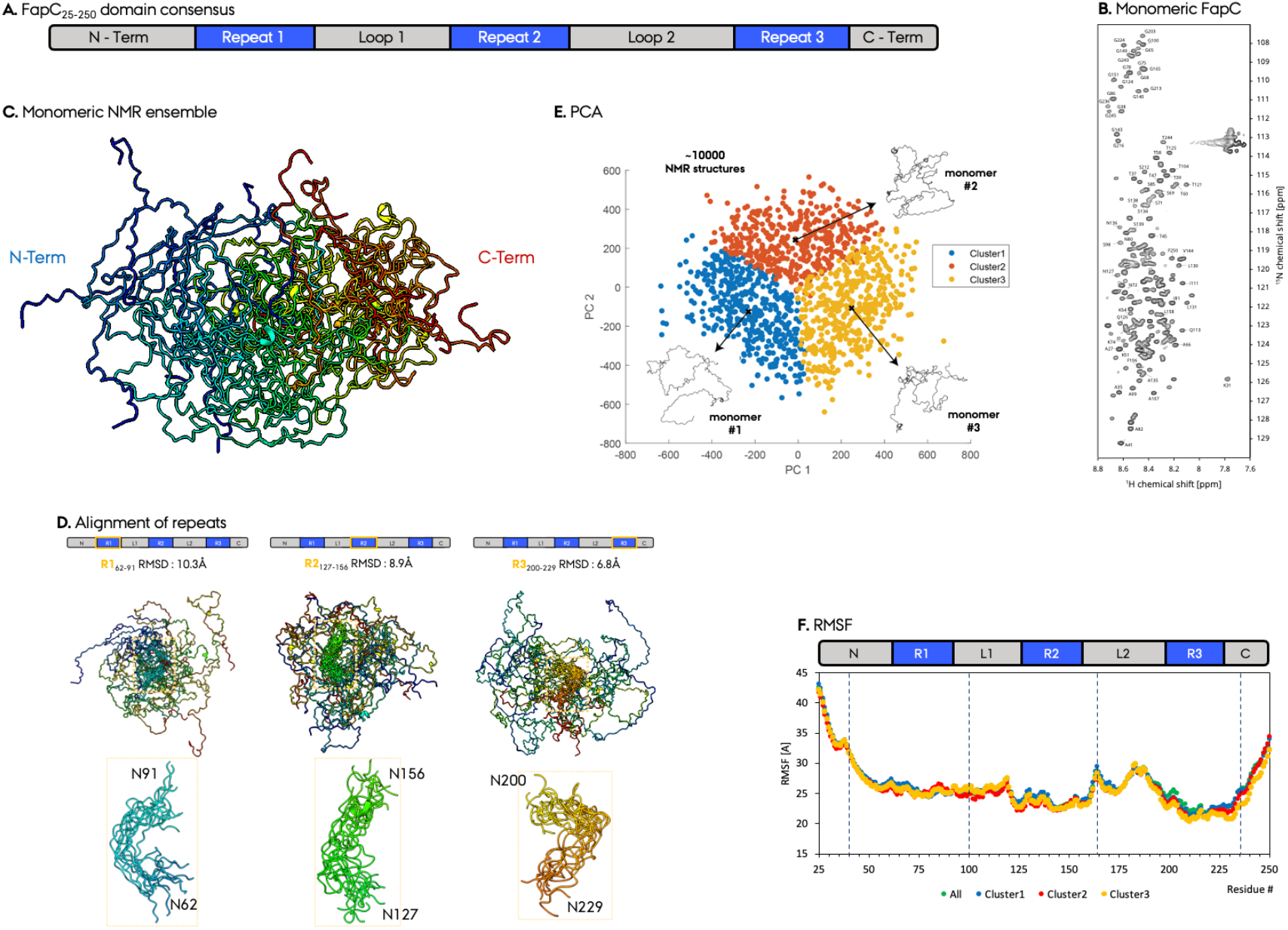
Solution NMR based ensemble description of monomeric unfolded FapC. **A**. Domain consensus of FapC functional amyloid. **B**. 2D ^1^H-^15^N HSCQ spectrum of FapC, with representative resonance assignments. **C**. FapC ensemble description from solution NMR. 10 minimum-energy structures are shown for representation, out of ∼10000 calculated structures. **D**. Alignment of three FapC repeats (R1-R3) for structures in C. **E**. PCA and **F**. RMSF analysis of FapC ensemble structures.

Using site-specific chemical shifts and nuclear Overhauser effect (NOE) distance restraints, we generated an extensive ensemble of ∼10,000 minimum-energy monomeric FapC structures, **Fig. SI2**. This large dataset allows unbiased analysis of FapC’s highly flexible monomeric state, a strategy successfully applied to other IDPs.^43^ The overlay of the 10 lowest-energy structures, aligned using the full-length sequence, **Fig. 1C**, shows an RMSD of ∼40 Å, reflecting the structural heterogeneity. Notably, the repeat regions (R1-R3) aligned with lower RMSD values (7-10 Å), suggesting a shared structural propensity among the repeats, **Fig. 1D**.

To further analyze structural variability, we performed PCA on the NMR ensemble.^42^ Projection onto the first two PCs followed by *k*-means clustering did not yield clearly separated conformational states. We divided the structures into three structural groups to select starting conformations for MD simulations, capturing the observed conformational heterogeneity (**Fig. 1E**). Residue-specific RMSF analysis quantified the flexibility of each region across the NMR ensemble (**Fig. 1F**) and revealed distinct flexibility patterns within the FapC domains: R3 exhibited the lowest RMSF values, followed by R2 and R1. These regions correspond to β-sheet-rich segments in both the Cryo-EM structure and the AF model (**Figs. 2, 3**). In contrast, the N- and C-terminal regions displayed the highest RMSF, consistent with their unstructured nature and high flexibility predicted by both Cryo-EM and AF.

**Figure 2:**
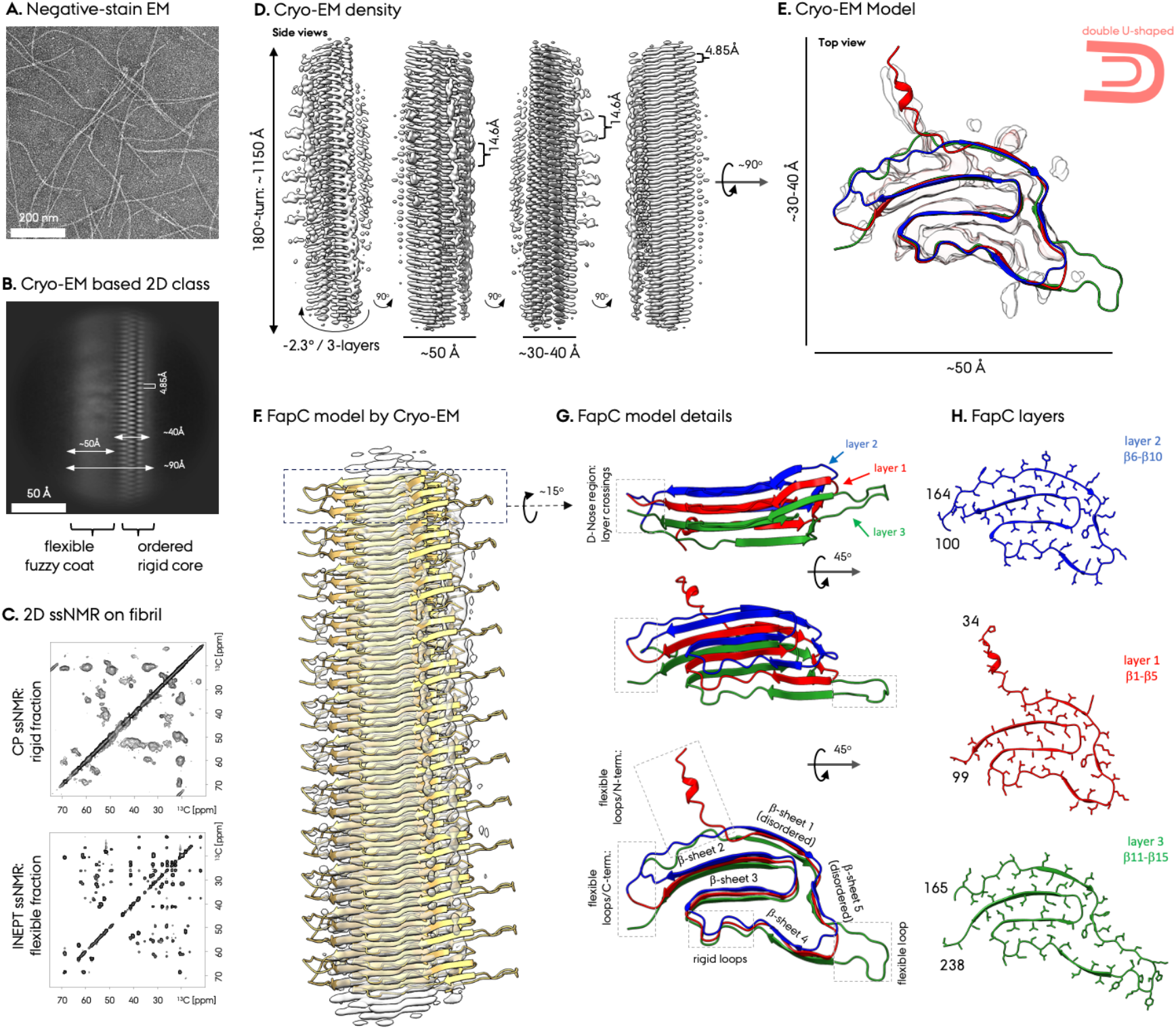
CryoEM characterization of fibrillar FapC. **A**. Negative-staining EM micrograph of FapC fibrils. **B**. CryoEM based reference-free 2D class average. The ordered fibril core and the fuzzy coat densities are indicative of rigid and flexible parts. **C**. 2D ^13^C-^13^C solid-state MAS NNR experiments probing rigid (via CP) and flexible (via INEPT) fractions of the FapC fibril. **D**. FapC CryoEM density map at high contour setting and four different orientations around the fiber elongation axis. **E**. One CryoEM FapC fibril subunit embedded in the CryoSPARC density map in red and the DeepEM enhanced map in light grey. **F**. The FapC model in the cryo-EM density. **G**. Details of the 3-layer structure of the FapC fibril subunit. Layers, layer-crossovers, β-sheets, and the flexible/rigid loop regions are indicated. **H**. Representation of the three unique layers within the subunit. The residue numbers representing the start/end of the layers are marked.

**Figure 3:**
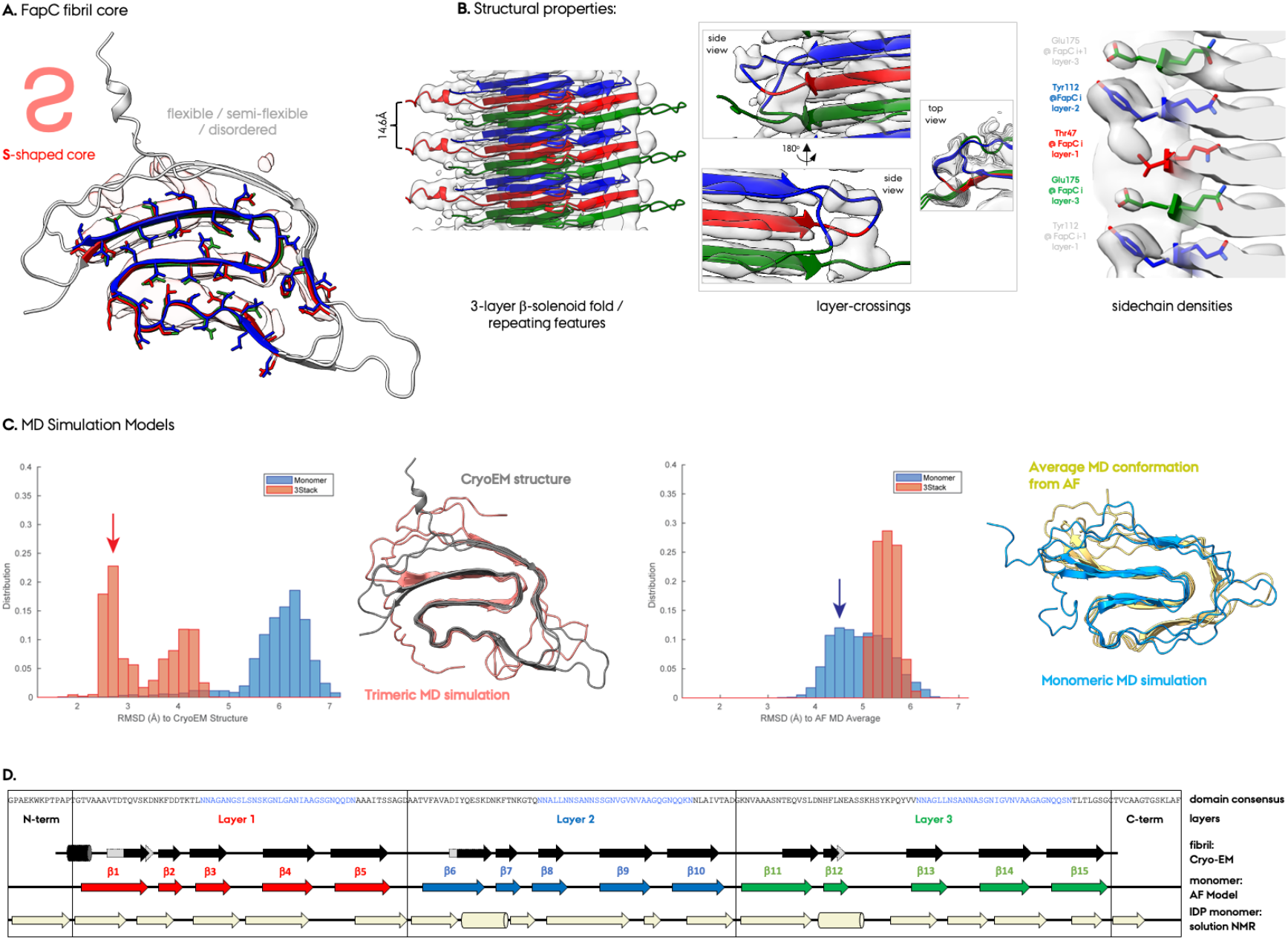
Structural properties of the FapC fibril from CryoEM and MD simulations. **A**. The core of FapC fibril has an S-shaped density. **B**. Structural features of FapC fibril. **C**. RMSD distributions of MD-sampled conformations of FapC performed as single and trimeric units, calculated relative to the CryoEM fibril model (first panel) and to the time-averaged structure obtained from MD simulations initiated from the AF model (third panel). RMSD values were calculated based exclusively on C_α_ atoms of the β strand regions observed in the CryoEM model. Blue and red arrows indicate the RMSD bins from which representative FapC conformations were selected for visualization. The second panel from left shows a representative FapC conformation from trimeric FapC MD simulations (red) superimposed onto the CryoEM structure (grey). The fourth panel illustrates a representative conformation of FapC from the single-unit FapC MD simulations (blue) superimposed onto the average structure derived from AF-initiated MD simulations (AF-MDav, yellow). **D**. Comparison of FapC secondary structure elements from solution NMR of unfolded monomer, AF monomer model and CryoEM fibril model.

### Folding Pathway from Unfolded FapC Monomer into Folded Monomeric Conformation

To identify the key FapC folding steps from unfolded IDP monomer into the folded monomer, we performed MD simulation on representative NMR determined monomers. For each of the three ensemble groups in the PCA, **Fig. 1E**, we selected the FapC structure that was closest to the arithmetic average of all structures as input for the MD simulations. We performed three separate 1000 ns MD runs for each of the three selected FapC monomers, yielding a total of nine simulations and 9000 ns of cumulative simulation time. We quantified the progression and extent of secondary structure formation in the MD simulations relative to the predicted AF folded monomer, **Fig. 3C**. Early in the folding process, the R3 repeat exhibited the highest propensity for β-sheet formation, reaching over 40% secondary structure content in three of the nine simulations by the end of each run. Similarly, R2 achieved more than 30% secondary structure content in four of the nine simulations. In contrast, R1 and L2 displayed lower β-sheet formation propensity (<20%), suggesting that R3 and R2 play a dominant role in the initial folding events, as well as the layers containing these repeat regions (Layer-3 with R3 and Layer-2 with R2). These findings correlate with previous mechanistic studies of FapC truncation mutants comprising different number of repeats, which showed that FapC lacking R2,R3 has the longest lag-phase in fibril formation.^13,15^ This supports our MD results and the critical role of R3 in FapC fibrilization, **Fig. SI4A**.

The correlation between higher secondary structure propensity in MD simulations and lower RMSF values in the NMR ensemble suggests that regions with reduced flexibility are more likely to adopt their native β-sheet rich conformations early in the folding process. This supports a sequestered folding mechanism, where β-strand-rich repeat regions serve as initial nucleation sites for the correct alignment of layers within a single FapC monomeric seed. Once this critical step is established, the subsequent folding of additional layers follows in a sequential manner, ultimately leading to the formation of the folded FapC monomer and its final fibril assembly.

### Unfolded monomeric FapC shares similar secondary structure with fibrillar FapC

To quantify the structural transition of FapC from its unfolded monomeric state to the fibrillar conformation, we performed ThT fluorescence assays and SR-CD spectroscopy. The ThT assay monitors fibril formation, as ThT selectively binds to β-sheet-rich fibrils, producing a fluorescence response, whereas both unfolded and folded monomeric FapC remain non-fluorescent, **Fig. SI4**. We observe an exponential curve without much lag phase in the aggregation kinetics for 50 μM FapC, **Fig. SI4**, in good agreement with previous FapC aggregation kinetics studies.^12,15^ In parallel, SR-CD spectroscopy was employed to assess the secondary structures of freshly desalted IDP unfolded FapC monomers and mature fibrils collected at the endpoint of the ThT fibrillization assay. Deconvolution of the SR-CD spectra reveals a shift from predominantly disordered monomers (29% β-strand, 8% α-helix and 63% loop/disordered) into fibrils with a predominantly β-sheet structure (40% β-strand, 10% α-helix and 50% loop/disordered). This is consistent with the minimum shifting from ∼200 nm in monomers to ∼220 nm in fibrils. Surprisingly, the unfolded monomers contain a relatively high amount of β-sheet propensity (30%) indicating structured elements in the unfolded monomers, correlating moderately with SSPs obtained by solution NMR (62% β-strand, 8% α-helix and 30% loop/disordered, with a maximum propensity of ∼20%).^19^ The higher β-sheet ratio could be due to initiation of aggregation in the samples during the preparation time following desalting, due to fast fibrilization of FapC. Comparative analysis of SR-CD data and NMR secondary structural characterization provides site-specific insights into the structural similarities of the unfolded monomer and fibrillar FapC. These findings suggest that specific secondary structure elements present in the unfolded monomer may serve as nucleation sites during fibril initiation and formation, guiding the transition towards the mature fibrillar state.

### Structure of Fibrillar FapC by CryoEM

Negative-stain micrographs of purified FapC fibrils revealed homogeneous filaments suitable for cryo-EM-based helical reconstruction and structure determination (**Fig. 2A**). 6969 movies were collected and the structure analysis was followed the CryoSPARC-based helical reconstruction approach we previously developed for the PSMα1 FA,^44^ **Fig. SI5**. High-quality, reference-free 2D-classification resulted in a single dominant class that showed clear layering of staggered β-strands, **Fig. 2B, SI5**. Unlike PSMα1, α-synuclein, and other amyloid fibrils,^44,45^ no evidence of polymorphism was observed in FapC fibrils.

The 2D class averages indicated that FapC fibrils are composed of a single protofilament with distinct high- and low-contrast regions. The high-contrast region corresponds to a rigid core ∼40 Å in width that is surrounded by the lower-contrast areas that corresponds to flexible loop, linker, and terminal regions, resulting in an overall fibril width of 80–90 Å, **Fig. SI6**. To further investigate structural heterogeneity and disorder, we recorded high-resolution 2D ^13^C-^13^C ssNMR spectra on isotope-labeled FapC fibrils, **Fig. 2C**. Overlaying spectra that selectively capture rigid (via CP) and flexible (via INEPT) regions revealed near-complete chemical shift coverage of amino acid types, **Fig. SI2**, confirming substantial dynamic differences within the fibrils, **Fig. 2E**, consistent with our previous findings.^15^

For 3D helical reconstruction, we refined the cryo-EM data without applying symmetry constraints. The helical parameters were determined to be ∼14.6 Å rise per FapC monomer (corresponding to three layers of ∼4.85 Å each) and a **−**2.3° helical twist per monomer, yielding a crossover distance of ∼1150 Å for a 180° turn comprising ∼80 FapC units and ∼240 layers (**Fig. 2D**). The final 3D density map revealed a well-resolved rigid core with continuous backbone connectivity and some visible side-chain density at an estimated resolution of ∼3.3 Å, **Fig. SI6**. The density map delineates three structural regions: (i) the rigid core with strong density, (ii) a semi-flexible part around the core with weaker density, and (iii) a fuzzy coat with no discernible structure with low density at low-contour levels, **Fig. SI6**,**7**. While most side-chain density was unresolved, the overall cryo-EM map provided sufficient resolution for structural modeling and refinement using the AF-predicted structure as the initial template. (**Fig. 2D-F, 3A, B**). The limited resolution of certain regions likely stems from flexibility and the unique three-layer fold of FapC fibrils, which present more challenges for subunit alignment not encountered in smaller, less flexible β-solenoid fibrils such as PSMα1 and CsgA or in planar amyloids like α-synuclein.^44,46,47^

Our previous solution NMR analysis and current SR-CD analysis of monomeric FapC indicate similar secondary structure propensities with the AF model,^19^ further supporting its validity. Using the cryo-EM density as a guide, we built the fibrillar FapC structural model by iteratively refining the initial AF model with ChimeraX ISOLDE MD to resolve steric clashes and unfavorable conformations. The final FapC fibrillar structure was validated using PHENIX,^37,38^ and exhibited excellent stereochemical quality with 97% of residues in favored Ramachandran regions with no outliers, along with low clash and MolProbity scores, **Table SI1**. This cryo-EM-based fibril structure comprised 48% β-strand, 2% α-helix and 59% loop/disordered regions. However, we note some ambiguity in low density regions due to intrinsic flexibility, which would slightly modify the secondary structure ratios, though these segments align well with the density at lower contour levels, **Fig. 2E, 3A, SI7**.

### Novel Structural Features of the FapC Fibril

The CryoEM structure of fibrillar FapC reveals a cross-β fibril with a distinct three-layer β-solenoid conformation, where flexible turns connect the layers, **Fig. 2G**. The β-strands are slightly tilted out of plane (∼15°) relative to the fibril axis, **Fig. 2F,G**, similar to the short PSMα1 fibrils but in contrast to the planar β-sheet arrangement observed in CsgA.^44,47^ The CryoEM density map, **Fig. 2F**, includes 12 FapC monomers that form a total of 36 stacked layers and accommodates residues 34–238. The termini (residues 25–33 and 239–250) are absent from the density, indicating high flexibility in these regions, consistent with the NMR results. Each three-layer subunit consists of residues 38–99 (Layer 1), 100–164 (Layer 2), and 165–236 (Layer 3). The fibril architecture comprises five β-strands per layer that collectively form five extended β-sheets spanning all three layers, **Fig. 2G,H**. A top-view reveals an interleaved double-U-shaped fold, **Fig. 2E,H**.

The CryoEM model of FapC shows several key differences with the AF monomeric fold, **Fig. 2G**, including curved and twisted β-strands that are instead straight in the AF prediction. The three central β-strands (β-strands #2–4, comprising residues 55–57, 63–96, 120–122, 128–161, 183–185 and 203–234) align closely with the AF model, forming an S-shaped rigid core with the strongest density, **Fig. 3A**,**SI7**. These strands define the most stable regions of the fibril. Conversely, β-strand #1 appears more disordered, deviating from the AF model, while β-strand #5 is notably shorter than predicted. Low-density regions in both the sharpened and the DeepEM-enhanced density maps are disordered and flexible, and correlate with the local resolution map and with low-confidence areas in the AF model, **Fig. SI7**.

The CryoEM model supports the three-layer β-solenoid fold proposed by AF, by clear repeating patterns in the CryoEM density that are separated by 14.6 Å, **Fig. 2D**. However, the stacking order within an individual FapC monomer appears irregular, following a 2-1-3 layer arrangement rather than the expected 1-2-3, **Fig. 3B**. The N-terminal region of each FapC monomer unit in the filament periodically extends out of the density every three layers, serving as a reference point for fibril alignment, **Fig. 3B**. Density at the crossover region suggests partial disorder within the three-layer fold, but still supports the overall stacking architecture. Three unique side chains that are on the outer surface, Tyr112, Thr47, and Glu175, are resolved in the CryoEM density, reinforcing the AF-predicted layer organization, **Fig. 3B**. However, given the current resolution, alternative layer arrangements cannot be entirely ruled out. The cryo-EM density confirms that FapC adopts a three-layer β-solenoid fold with an S-shaped rigid core featuring curved β-strands, disordered regions, and an irregular layer stacking pattern. While the structure aligns with the AF model in many aspects, deviations in strand curvature, layer order, and flexible regions highlight the unique dynamic nature of FapC fibrils comprising both rigid and flexible areas.

### Oligomeric Interfaces are Crucial for the Fibrillar FapC Fold

AF predicts the structure of FapC based primarily on its amino acid sequence, but is not yet well suited to predict amyloid fibril structures due to methodological limitations and available training data.^48^ In contrast, the CryoEM map is obtained from FapC fibrils where each FapC subunit makes extensive intra- and intermolecular contacts. Consequently, the CryoEM structural model differs from the AF-predicted monomer model by reflecting the impact of the oligomeric environment. The RMSD between the two full-length structures (all C_α_ atoms, residues 34–238) is 6.8 Å, indicating substantial differences; however, this value is strongly influenced by flexible loop regions where AF predictions show high structural uncertainty, **Fig. 3**. In comparison, the structured regions show less divergence in the overall β-strand conformation (RMSD: 5.2 Å), particularly in the S-shaped core region (RMSD: 4.6 Å), **Fig. SI7**.

To explore the origin of structural differences between the cryo-EM fibril structure and the AF monomer model, we performed three independent 1 μs-long MD simulation runs, initiated from (1) the AF monomer model, as well as three 400 ns-long MD simulation runs each initiated from; (2) one subunit of the cryo-EM-derived fibril structure; and (3) a trimer from the CryoEM model to retain the packing effects in the filament. To quantify the structural similarity, we compared the RMSD of the β-sheet regions in the cryo-EM model with the structures from the MD simulations, **Fig. 3C**. Simulations initiated from the AF monomer model (1) were averaged (termed AF-MDav) and maintained a moderate RMSD of ∼5.6 Å for all C_α_ atoms and ∼1.5 Å for the β-strand C_α_ atoms compared to the initial AF model. The low-confidence regions of the AF model, which predominantly correspond to unstructured regions, deviated significantly from their initial conformations and exhibited high flexibility during the MD simulations. In contrast, the β-strand conformations remain largely stable. AF-MDav, **Fig 3C**, serves as a reference to determine whether the cryo-EM FapC structure would deviate from its fibrillar fold and shift towards an AF-like conformation, or instead remains close to its original state. Simulations initiated from the CryoEM FapC structure as a single unit (2) diverged significantly from the initial structure, showing a clear tendency to deform towards the AF conformation (β strand RMSD values, corresponding to distribution peaks in Fig. 3, were 6.3 Å relative to the initial cryo-EM structure and 4.5 Å relative to AF-MDav). This suggests that without intermolecular contacts, an isolated FapC monomer tends to adopt a more AF-like conformation. To assess the role of subunit-subunit interactions, we simulated the cryo-EM FapC structure in a trimeric arrangement (3) as a control condition, **Fig. 3**. The top and bottom FapC units were constrained at C_α_ atoms, while the central subunit remained free to move during the MD simulations. Strikingly, this unconstrained central subunit retained its cryo-EM fold, exhibiting a β-strand RMSD distribution peak of 2.7 Å relative to the cryo-EM structure and did not shift toward the AF-MDav average conformation (β-strand RMSD of 5.5 Å), **Fig. 3**.

These MD results suggest that the earliest folded monomeric FapC conformations differ from those found in the mature fibrillar state, yet the core structure remains largely similar. We propose that the MD-optimized monomeric structures represent a nascent FapC fold, which undergoes subtle conformational changes driven by protein-protein interactions within the fibril to reach the mature fibrillar (seed or seeding-competent) conformation. This seed or seed-like state subsequently assembles into fibrils via elongation. However, the precise mechanism underlying this conformational transition remains to be elucidated.

### Key Transition Steps in FapC Folding and Unfolding

To investigate the key stages of the folding mechanism, we performed SMD simulations to systematically unfold the folded FapC structure, assuming that folding occurs in the reverse order of unfolding. Using the AF predicted monomeric fold, the initial monomeric FapC conformations were extracted from equilibrated MD trajectories at time points 100, 200, 300, and 400 ns. These served as starting structures for the SMD simulations. The choice of the AF folded monomeric model was based on its relevance to the early folded FapC conformation, which is expected to transition into the seeding-competent fibrillar FapC fold. In total, 12 separate 700 ns-long SMD simulations were conducted (totaling to 8,400 ns). To identify key structural transition events, we monitored inter-β-sheet and inter-layer distances, which provide insight into the “unfolding” process as a potential reverse-folding mechanism (**Fig. 4**). Since FapC adopts a unique 3-layer fold, it was essential to analyze unfolding in three dimensions. Inter-β-sheet distances captured conformational changes in the xy-plane, and inter-layer distances tracked unfolding along the z-axis (fiber axis). Based on the distribution of rupture or separation events in inter-β-sheet and inter-layer contacts, the most likely unfolding pathway proceeds through five steps: i) sheets 1 and 2 separate; ii) layers 1 and 2 dissociates within sheet 1, while sheets 2 and 3 remain intact; iii) sheets 3 and 4 separate; iv) layers 1-3 dissociate; and v) finally sheets 3 and 4 detach completely, leading to a fully unfolded state that has lost all tertiary structure, **Fig.4, SI3**.

**Figure 4:**
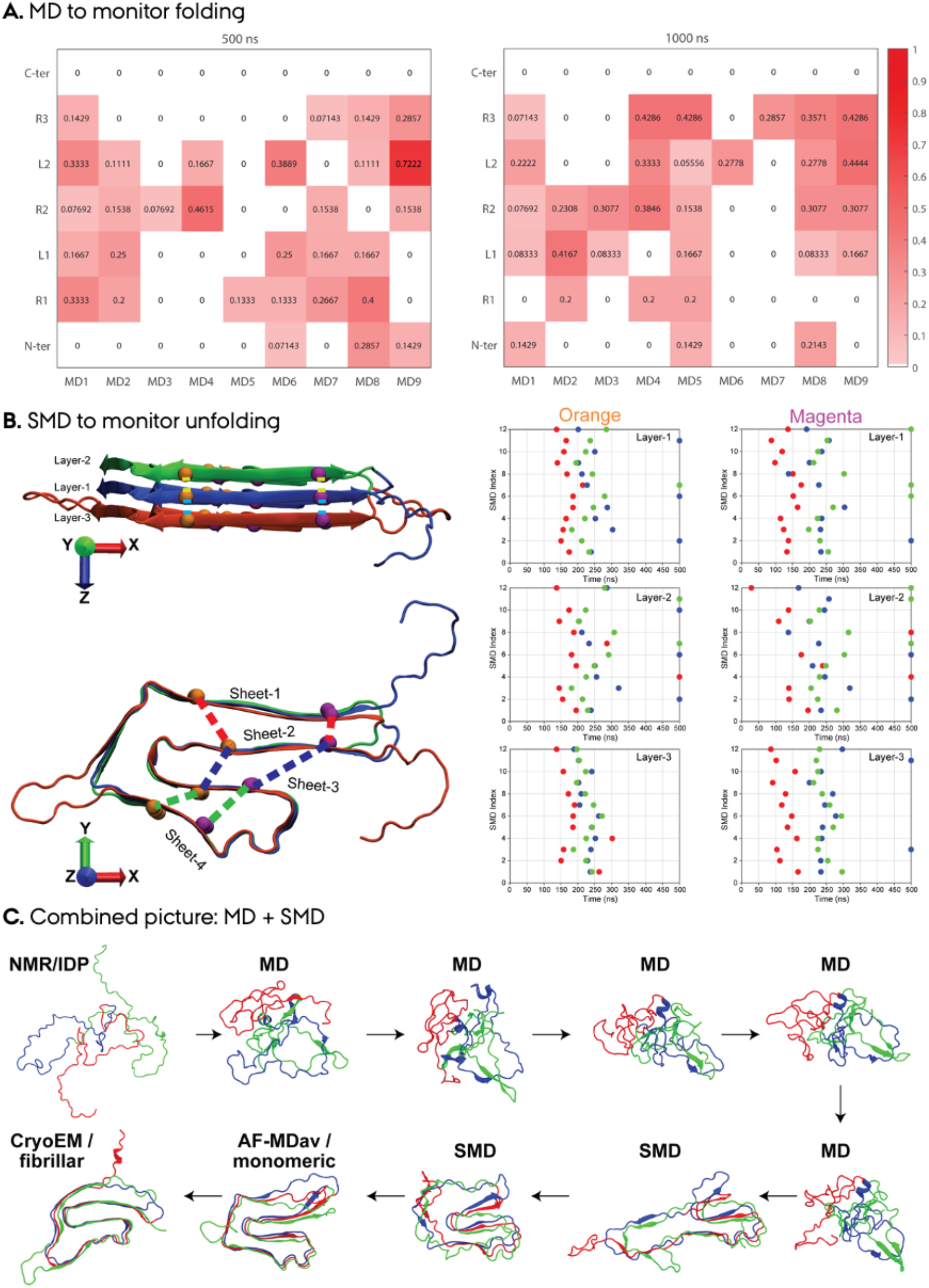
MD simulations tracking the transition from unfolded monomer to fibril. **A**. MD simulations were initiated from the monomeric unfolded FapC NMR structures, shown in Fig. 1E. The MD simulations were run for 1 µs and the changes in the secondary structure formation monitored. The heatmap shows the percentage of secondary structure formed in each FapC consensus structural domains (N-term, R1, L1, R2, L2, R3, and C-term) for each individual run relative to the AF-MDav. The left heatmap represents structures at 500 ns, while the right heatmap reflects the end of the simulations at 1000 ns. **B**. To probe the FapC unfolding mechanism and pathway, we performed SMD simulations, which were initiated from MD-equilibrated AF models, and monitored the changes in key distances that determine FapC fibril fold, for three layers and four β-sheets. The schematic representations (left) illustrate protein conformations with dashed lines indicating measured distances between selected C_α_ atoms (highlighted by orange and magenta beads). These distances were chosen to track the dissociation of fibril-like interactions between β-sheets and layers. The dot plots (right panels, titled “Orange” and “Magenta”) indicate simulation time instances when these distances surpass a 12 Å threshold, signaling significant structural rearrangements or separation events. Those that did not reach the 12 Å threshold within the simulation time are shown at the 500 ns time point. Dots in these plots are color-coded consistent with the dashed lines in the schematic representations, clearly linking each measured distance to its corresponding position within the protein structure. Layers 1, 2, and 3 are depicted in blue, green, and red, respectively. **C**. Selected conformations from MD simulations (MD9, initiated from NMR structures) and SMD simulations (SMD2) illustrate the progression from the unfolded NMR structure through partially folded intermediate states, to a folded monomeric state, and ultimately to a fully folded fibrillar structure. For the depiction of the SMD simulations and the final SMD conformation, please refer to Fig. SI3.

When combined with our MD simulations initiated from NMR-based unfolded FapC monomer, which monitored the initial stages of the folding pathways, the SMD simulations provide detailed molecular insights into the complete folding/unfolding mechanism of FapC (**Fig. 4)**. Our simulations suggest that secondary structure formation (within individual β-sheets and layers) precedes tertiary organization (i.e., the correct arrangement of layers in the final 3-1-2 order). However, this does not rule out the possibility that some tertiary contacts may form concurrently or even before all secondary structural elements have fully matured.

## 4. Discussion: From Monomeric Disorder to Functional Amyloid Fibril

We have applied an integrative structural biology approach to characterize the diverse states of the functional amyloid FapC from Gram-negative *P. aeruginosa* that presents a holistic picture of amyloid fibril formation in the context of biofilm-forming FAs. The three distinct states (unfolded monomer, folded monomer, and fibril) illustrate the dynamic biogenesis of FapC involving significant structural reorganization and conformational changes and highlights key regulatory steps and potential intervention points.

The large structural ensemble generated using our previously determined chemical shift assignments together with the NOESY restraints reported above provide a robust framework for evaluating structural features of the unfolded FapC monomer. Although adopting a largely random coil conformation, the unfolded monomeric FapC exhibits a significant SSP of ∼20% that follows closely the AF folded monomer model,^19^ suggesting a smooth transition from the IDP state to the folded monomeric conformation. Our MD-optimized AF model, and consequently also AF, represents this folded FapC monomer as an intermediate stage in fibrillization that will form the final fibrillar FapC fold determined by cryo-EM upon conformational changes. The overall secondary structure composition remains relatively constant throughout these transitions between the unfolded monomer (despite its low SSP), the AF monomer model, and the fibrillar cryo-EM structure, **Fig. 3G**. SR-CD spectroscopy supports these structural assignments and also indicates a greater proportion of ß-sheet conformations in the monomeric FapC (8% α-helix, 29% β-sheet, 63% disordered/turns) compared to fibrils (10% α-helix, 40% β-sheet, 50% disordered/turns), **Fig. S4**. These SR-CD measurements may reflect a mixed state comprising both unfolded and folded/semi-folded monomer. Notably, solution NMR-derived SSPs correlate with these CD-based ratios.

The cryo-EM fibril structure alongside the AF monomer model reveal domain arrangements distinct from the NR1L1R2L2R3C consensus for FapC_25-250_.^9,21^ The original domain arrangement is based on sequence determination of the three imperfect repeats and does not take the structural elements into account. Our structural model expands upon the previously defined repeat regions (R1-R3), demonstrating that the 3-layer fibril fold not only incorporates these repeats but also utilizes the previously designated loop regions as integral structural elements. While our prior work demonstrated that truncating these loops affects fibrillization, the new cryo-EM fibril model provides the structural basis for this observation, showing that loops L1 and L2 form β-sheet regions that contribute directly to the fibril core. This model explains why aggregation has previously been observed for FapC even after deletion of R1, R2 and R3.^13^ The observed 3-layer architecture significantly extends the repeat regions while minimizing loop flexibility, **Fig. SI1**, leading to a compact and stable fibril structure.

PCA analysis of unfolded FapC monomer ensembles confirmed the absence of distinct subpopulations, while RMSF analysis highlighted the structural alignment of R2 and R3 with lowest scores. The N- and C-terminal regions, as well as the longer loop regions (L1 and L2), remain highly flexible. MD simulations (9 µs in total) of representative IDP monomer structures revealed a stepwise transition pathway to the folded monomeric state. Notably, the R3 region exhibited the highest β-sheet propensity (>40% in 3/9 simulations) followed by R2 (>30% in 4/9 simulations), consistent with the lower RMSF values observed in the NMR ensemble. In contrast, R1 and L2 showed minimal secondary structure formation. These data suggest that the unfolded-to-folded monomer transition is initiated by the key R3 region which serves as a nucleation interface, followed by the stepwise formation of intramolecular β-strands and then layers. This model is in good agreement with our previous observations that removal of R3 and R2,R3 result in extended lag-phase compared to full-length FapC and removal of other repeats.^13^

Cryo-EM analysis revealed that FapC forms a homogenous fibril composed of a single protofilament with a ∼40 Å diameter rigid core featuring well-ordered β-strand stacking, surrounded by a ∼50 Å flexible “fuzzy coat.” The estimated resolution of ∼3.3 Å aligns with previously reported biofilm-forming FA fibril structures, including CsgA (up to ∼4 Å by cryo-EM), PSMα1 (∼3.4 Å by cryo-EM), TasA (1.6/∼3.5 Å by X-ray crystallography and cryo-EM), and PSMα3 (1.5 Å by X-ray crystallography).^44,47,49-52^ Notably, native FapC filaments extracted from *P. aeruginosa* biofilms have an average width of 8 ± 2 nm, similar to the *in vitro* assembled FapC fibrils, as observed previously for CsgA and TasA, **Fig. SI4**.^47,50^ The *in vitro* filaments are longer compared to the *in vivo* filaments likely due to extensive processing of the *in vivo* fibrils during the extraction protocol.

Our all-atom MD simulations of different FapC structural states underscore the role of oligomeric interfaces in stabilizing fibril architecture. While the AF folded monomer model remained relatively stable, the single cryo-EM fibril subunit exhibited significant deviations, shifting toward an AF-like structure. However, the trimeric cryo-EM fibril model maintained structural integrity, emphasizing the importance of subunit-subunit interactions in fibril stabilization.

The fuzzy coat constitutes nearly half of the cryo-EM density, consistent with MD simulations and low-confidence regions in AF predictions that highlight structural disorder and flexibility, **Fig. SI6**,**7**, unlike other biofilm-forming FAs such as CsgA, TasA, and PSMα1. This suggests a unique role of FapC in biofilm matrix formation and interactions. The helical twist of FapC fibril is small due to its 3-layer architecture, reducing apparent twisting three-fold, with a -2.3° twist and rise of ∼14.6 Å per monomer. While the handedness remains undetermined, we assumed a left-handed filament, consistent with common amyloid structures. The fibril adopts a double-U-shaped architecture with an S-shaped core, aligning broadly with AF predictions but deviating significantly due to oligomeric interface interactions and differential flexibility. Whether the FapC fibril undergoes conformational maturation from a preliminary fold described by the AF folded monomer model to its final cryoEM fibril fold remains an open question, but this has been observed for α-synuclein and hIAPP fibrils.^45,53,54^

Unlike the pronounced polymorphism observed in pathological amyloids such as α-synuclein, larger biofilm-forming FAs like CsgA and TasA exhibit remarkable structural homogeneity, both in vitro and in vivo.^47,50^ However, our recent structural analysis of FA PSMα1, a small 21 amino acid peptide, showed that it is polymorphic with six different folds.^44^ In contrast, our cryo-EM data of FapC reveal a single fibril class, suggesting a highly uniform assembly process, even in the absence of its *in vivo* partners, FapB and FapE, **Fig. SI5**,**9**. AF modeling suggests that FapB shares structural similarity with the FapC fibril core, particularly in β-strands 2-3, while exhibiting greater disorder in β-strands 1 and 5, **Fig. SI9**. This aligns with the flexible regions identified in MD simulations and supports the hypothesis that FapB templates FapC fibril formation. Overall, these support the consensus that FapC fibril is partially disordered and templated by FapB having a similar fold and gives structural insights into Fap biogenesis. Despite their functional differences, β-solenoid fibrils share common inter-strand hydrogen bonding and hydrophobic core packing to maintain fibril integrity, **Fig. SI9**. Het-S and CsgA adopt a well-ordered β-solenoid fold with tightly packed layers, whereas FapC and HELLF exhibit increased structural flexibility due to longer loop regions extending from the fibril core.^47,55,56^ Canonical ordered stacking of layers is observed in all of these amyloids (layer 1-2-3 repeated). However, our cryo-EM density map doesn’t rule out the irregular three-layer β-solenoid fold (layer 3-1-2) of the AF folded monomer model, **Fig. 3B**, where layer 1 is sandwiched between layers 2 and 3. Such an organization would suggest a sequestered FapC folding pathway facilitated by flexible loop regions that guide correct layer positioning. The presence of extended loop and turn regions in FapC enables this irregular three-layer fold. MD and SMD simulations support this hypothesis, providing insights into the sequential order of layer formation.

The electrostatic potential map of FapC filaments reveals charge distribution patterns that may contribute to fibril stability, **Fig SI6**. Hydrophobic interactions play a key role in assembly, stabilizing inter-sheet contacts and facilitating monomer incorporation. These structural, electrostatic and hydrophobic properties likely influence *Pseudomonas* biofilm formation and interactions of FapC with eDNA and polysaccharides.

Finally, our results suggest that the transition from an unfolded monomer to fibrillar FapC proceeds through multiple structural states, with key intermediates likely mirroring its assembly and disassembly pathways, **Fig. 5. State-1:** unfolded monomer with preformed secondary structure elements. **State-2:** intermediate folded monomeric conformation forms a stable cross-β structure through small-scale structural rearrangements. **State-3:** small conformation changes adjust the folded monomer to the fibrillar fold. **State-4:** cross-β FapC mature filaments form by a stable cross-β network. The fibril disassembly pathway likely mirrors its assembly process, with specific conformational states serving as key intermediates.

**Figure 5:**
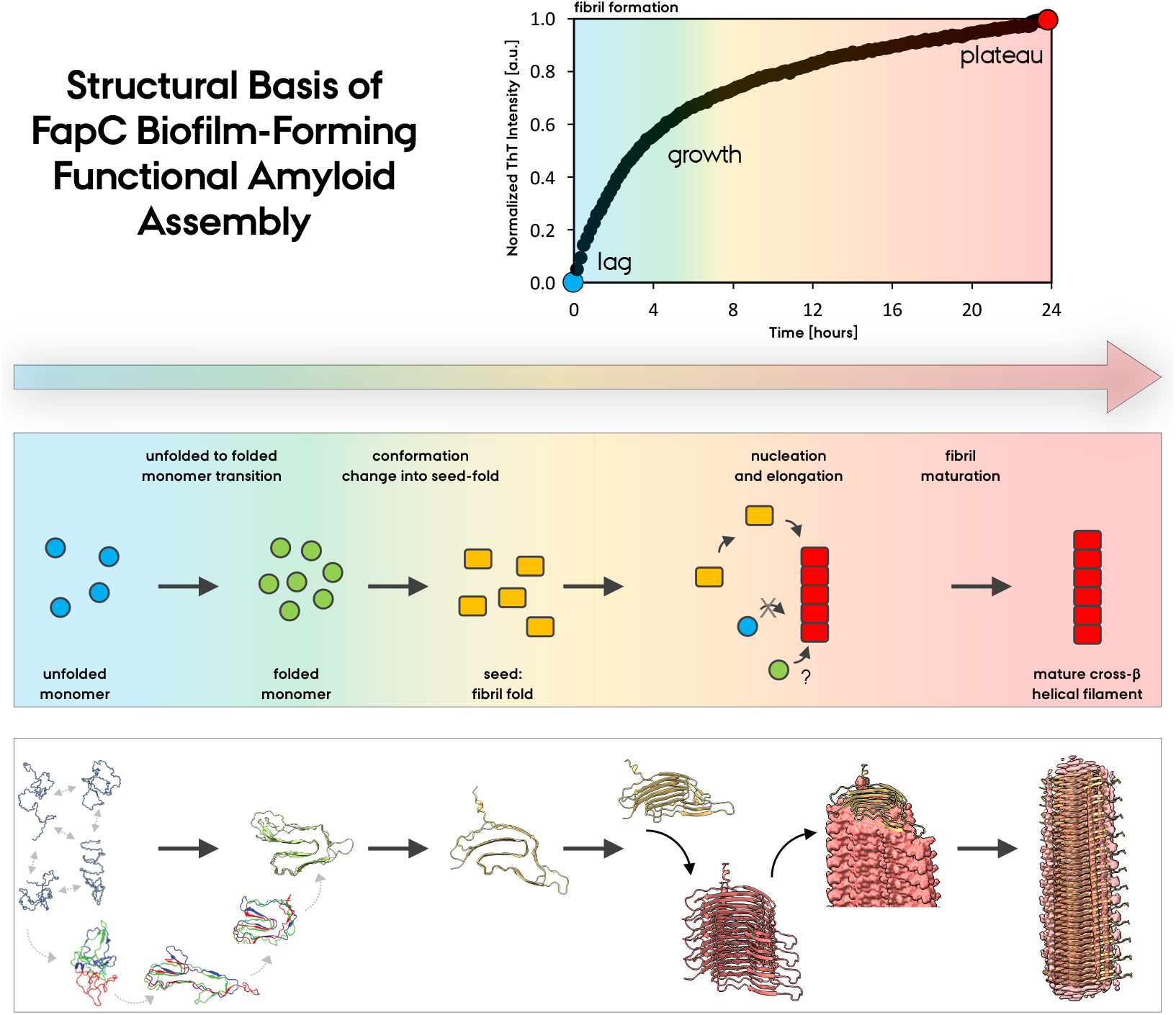
Structural model for FapC biofilm-forming functional amyloid formation. Combining solution NMR, MD/SMD simulations and CryoEM, we present the structural basis of FapC functional amyloid fibril formation in *Pseudomonas* comprising monomer to fibril structural transition via a folded monomeric intermediate. The top panel shows the ThT assay used to monitor fibrilization formation. Middle panel describes the key steps. The bottom panel describes the remarkable structural transition in FapC biogenesis: from unfolded monomer, to folded monomer and finally to fibrillar FapC.

## 5. Conclusion

The fibrilization initiation model we present here provides a complete structural and mechanistic understanding of FapC assembly and the biogenesis of functional amyloid in pathogenic *P. aeruginosa* biofilm formation. This information serves as a foundation for developing anti-biofilm and anti-amyloid therapeutics to combat biofilm-associated antimicrobial resistance. The strong structural correlation between AlphaFold predictions and experimental models, along with the ability of MD simulations to refine and bridge structural gaps through biophysically meaningful refinements, highlights the predictive power of computational approaches in amyloid research. The distinct organizational features of FapC suggest a specialized assembly mechanism that differs from other known functional and pathologic amyloids. Understanding these structural and functional characteristics not only deepens our knowledge of bacterial biofilms but also informs broader amyloid research, including potential applications in biotechnology and disease-related amyloid studies.

## Data Availability

All data can be requested from the corresponding authors. The FapC fibril density map has been deposited in the EM Databank with accession number EMD-**49649** and the atomic model in the Protein Databank with accession number **9NQD**.

## Author Contribution

Conceptualization: UA; Methodology: MG, UA; Data collection, formal analysis and investigation: KH, MGo, CHB, AT, EBP, JFC, MA, MG, UA; Writing - original draft preparation: All; Funding acquisition: UA; Resources: MG, UA; Supervision: MG, UA. All authors reviewed the manuscript.

## Acknowledgements

UA acknowledge support from the Department of Structural Biology, University of Pittsburgh School of Medicine (UPSOM) for access to the high-field NMR and Cryo-EM facilities. The Pittsburgh Center for CryoEM (RRID:SCR_025216) was supported in part by the UPSOM, the Department of Structural Biology, and the National Institutes of Health (grants #S10-OD-019995 and #S10-OD-025009). The content is solely the responsibility of the authors and does not necessarily represent the official views of the National Institutes of Health. We acknowledge the beam time on the AU-CD beam line at ASTRID2. We thank Dr. Ulasli and Dr. Byeon for their help in EM and solution NMR. UA acknowledges start-up funding by the UPSOM, and a Competitive Medical Research Fund grant by the UPMC Health System. MA acknowledges financial support from Aarhus University Research foundation NOVA (AUFF-E-2023-9-18), Magda Sofie, Aase Lütz’s Memorial Trust and Hørslevfonden.

