## Supplementary material for "Structural Basis of *Pseudomonas* Biofilm-Forming Functional Amyloid FapC Formation": Supplamentary Tables

### Supplementary Data: Tables

**Table SI1: Cryo EM summary:** Summary of cryo-EM data collection, reconstruction parameters, model building, and structure validation scores by Phenix.

| **Data collection** | |
|---|---|
| Microscope | Titan Krios G3i |
| Energy filter | Selectris, 10eV slit |
| Camera | Falcon 4i |
| Voltage | 300 kV |
| Magnification | 165,000x |
| Defocus range | -0.5 to -2.0 µm |
| Pixel size | 0.72 Å |
| Exposure time (s) | 4.8 s |
| Electron dose | 45 e/Å$^2$ |
| **Reconstruction** | **CryoSPARC** |
| Box size | 300-pixel |
| Separation distance | 84.6 Å |
| Number of initial particles | ~762,000 |
| Number of final particles | ~72,000 |
| Symmetry imposed | C1 (None) |
| Helical rise/helical twist | 14.562 Å / -2.266° |
| Map resolution @0.143 FSC | 3.3 Å |
| **Structure validation statistics** | **PHENIX** |
| MolProbity score | 1.44 |
| Clash score | 4.47 |
| Ramachandran favored | 97 |
| Ramachandran allowed | 3 |
| Ramachandran disfavored | 0 |
| RMSD of bonds | 0.015 Å |
| RMSD of angles | 2.660 |
| Number of chains in the model | 12 |

**Table SI2: Updated selections of Fixed and pulling atoms in SMD simulations after 100Å extension.** The new fixed and pulling atoms were set to the N- and C-termini of the structured fold, enabling continued simulation of the mechanical unfolding behavior for the remaining structured regions in a controlled manner.

| | Fixed Atoms | SMD Atoms |
|---|---|---|
| SMD1 | T39, L61, Q126 | L233, N225, L205 |
| SMD2 | A42, S50, F55, Q126 | T230, A220, G203, F183 |
| SMD3 | N53, N62, Y112 | Q227, I215, V146 |
| SMD4 | V49, A82, N122 | T232, K190, Q150 |
| SMD5 | V40, T47, K54, Q126 | G234, S228, F183, L204 |
| SMD6 | T39, L61, I81 | L233, S228, L204 |
| SMD7 | P36, N53, A83 | S228, V217, G203 |
| SMD8 | P36, N53, N80 | S228, I215, H182 |
| SMD9 | G38, T47, S85 | L231, N225, G203 |
| SMD10 | T39, F55, I81 | L233, V217, N201 |
| SMD11 | V49, A83, I111 | G234, I215, N181 |
| SMD12 | V40, N62, Y112 | T232, Q227, G203 |
